# RhoGEF12 regulates endosomal SORL1-retromer and its inhibition is therapeutic in human neuronal models of Alzheimer’s disease

**DOI:** 10.64898/2026.03.06.709427

**Authors:** Yasir H. Qureshi, Charles A. Williams, Istvan Hajdu, Suvarnambiga Kannan, Ananya Govindarajan, Barbara Végh, Gregory A. Petsko, Jessica E. Young, Péter Závodszky, Scott A. Small

**Author notes:** Contributed equally (Yasir H. Qureshi, Charles A. Williams).

## Abstract

The interaction of the endosomal sorting protein SORL1 with the retromer complex at endosomal membranes controls a recycling pathway whose dysfunction is pathogenic in Alzheimer’s disease (AD) and is linked to other neurodegenerative disorders. To search for novel therapeutic targets, we hypothesize that endosomal SORL1-retromer might be regulated by SORL1’s cytoplasmic tail. We begin by completing an *in vitro* analysis of the tail and show that its phosphorylation by ROCK2 (Rho-associated kinase 2) reduces SORL1’s affinity to retromer. Since RhoGEF12 (Rho guanine nucleotide exchange factor 12) is an upstream activator of ROCK2 that is upregulated in AD, we used a RhoGEF12 pharmacological inhibitor to mechanistically and therapeutically validate the findings in neuronal cultures. First, in mouse neurons we confirm that the inhibitor increases endosomal SORL1-retromer. Next, we turned to human iPSC-derived neurons to show that the inhibitor reduces Aβ40 and Aβ42, an indicator of pathway upregulation, in a SORL1-dependent manner. Finally, we validate its therapeutic potential by applying the RhoGEF12 inhibitor to human iPSC-derived neurons expressing AD-associated mutations in either *APP* or *SORL1*. Collectively, our results identify a novel and therapeutically amenable mechanism that regulates endosomal SORL1-retromer and preclinically validate RhoGEF12 as a therapeutic target for AD and potentially other neurodegenerative disorders.

**One sentence summary:** Pharmacological inhibition of RhoGEF12 increases endosomal SORL1-retromer recycling and reduces pathogenic amyloid secretion in human neuronal models, identifying a novel, targetable pathway for treating Alzheimer’s disease.

## INTRODUCTION

Retromer is an evolutionarily conserved protein assembly that functions as a "master conductor" of intracellular trafficking^1^. It primarily operates on endosomes to sort and rescue specific transmembrane proteins from being sent to the lysosome for degradation, instead recycling them back to the plasma membrane or the trans-Golgi network. The heterotrimer comprising VPS26-VPS35-VPS29 is retromer’s core complex that, among other functions, acts to coat and stabilize endosomal recycling tubules^2^.

The type I transmembrane protein SORL1 (‘sortilin-related receptor 1’, also called SORLA or LR11), is highly expressed in the brain^3^. While expressed in all brain cell types, the protein is preferentially found in neurons^4–8^. A convergence of genetic, molecular, and cellular studies have established that SORL1’s interaction with the retromer complex at endosomal membranes represents a functional unit that can control endosomal recycling, and that pathway disruptions are pathogenic in Alzheimer’s disease (AD)^9^. The pathway is neuroprotective in part by controlling the health of lysosomes and autophagosomes^10^, which can account for why the pathway is also implicated in other common neurodegenerative disorders^11^. Together with studies establishing its pathogenicity in Parkinson’s disease (PD)^12–15^, more recent studies have linked pathway disruptions to primary tauopathies^16^, TDP-43 frontotemporal degeneration (TDP-43-FTD)^17^, and amyotrophic lateral sclerosis (ALS)^18,19^.

There is an urgent need, therefore, for interventions that upregulate SORL1-retromer endosomal recycling. We have previously introduced a class of small molecules that were designed to bind and stabilize the retromer complex^20^. While they have been shown to be useful tool compounds in boosting the pathway in model systems of AD^8,20–22^, ALS^19^, and PD^23,24^, because of their low potency and specificity they do not appear suitable as human drugs. We have thus turned our attention to SORL1, informed by a recent study showing that a genetic manipulation that increases endosomal SORL1-retromer enhances pathway function by reducing the production of both Aβ40 and Aβ42^11^. This readout has emerged as one of the most reliable biomarkers of pathway function, because SORL1-retromer recycles the amyloid precursor protein (APP) out of endosomes, an organelle where APP is proteolytically cleaved to produce these amyloid peptides^11^.

In considering novel interventions that might increase endosomal SORL1-retromer, we were guided by a proteomic screening study that found that ROCK2 (Rho-associated kinase 2) is one of the top SORL1 binding partners^25^, and from studies suggesting that the SORL1’s 54-residue cytoplasmic tail contains putative ROCK2 phosphorylation sites^25,26^.

Mechanistically relevant, the tail has three established binding motifs that account for why SORL1 highly localizes to early, retromer-positive, endosomal membranes^27–29^. As illustrated in Figure 1, the first two allow SORL1 to bind the endosomal sorting proteins GGA1 and AP1^30^. This binding ensures that, when at the trans-Golgi network, SORL1, like a handful of other transmembrane proteins containing similar motifs, is actively transported to endosomes. The tail’s third binding motif, one unique to SORL1, is to the retromer protein VPS26^28^. VPS26 binding dictates that when arriving in endosomal membranes, SORL1 gets incorporated into the retromer heterotrimeric complex. As also illustrated in Figure 1, SORL1 is found to dimerize via its 3FN domain in retromer-positive endosomes, and recent studies have begun modelling SORL1’s folded structure^27^.

**Figure 1.**
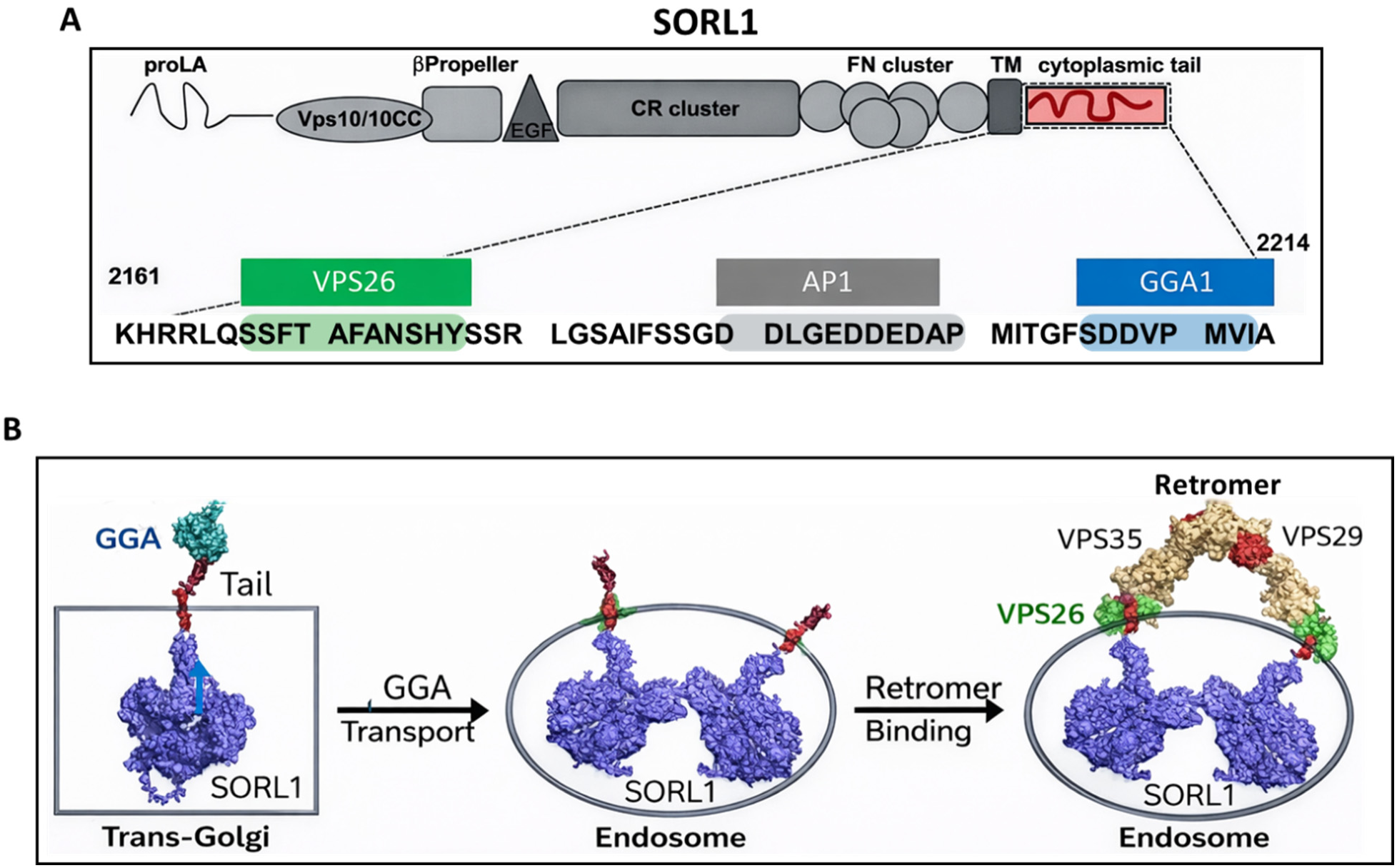
SORL1’s cytoplasmic tail regulates endosomal SORL1-Retromer. **A.** SORL1’s multiple domains are illustrated with its cytoplasmic tail enlarged to highlight its GGA1, AP1, and VPS26 binding motifs. **B.** An illustration showing that, by binding GGA1 at the trans-Golgi network, SORL1 is actively transported to endosomes, where it dimerizes. By VPS26 binding, the SORL1-retromer functional unit is formed, which regulates an endosomal recycling pathway implicated in Alzheimer’s disease and other neurodegenerative disorders. Although the structure of the retromer heterotrimeric core, with VPS35 in beige, VPS29 in red, and VPS26 in green, is taken from an actual cryoET structure by the Collins group (2), the structure of the SORL1 dimer is only a hypothetical model.

Besides compacting this long protein and enabling it to fit inside endosomes, the dimerization step has been shown to be critical for retromer binding and pathway function^27^, since it aligns folded SORL1 inside the endosome with retromer dimers on the endosome surface.

An *in vitro* analysis is important to definitively establish precisely where ROCK2 interacts with the tail and for understanding its functional consequence. We therefore begin by synthesizing tail peptides *in vitro* and find that ROCK2-dependent phosphorylation is highest at the tail’s N-terminal end, and that this phosphorylation dramatically reduces VPS26’s binding affinity to the tail. Since ROCK2’s phosphorylation ability only occurs in its activated state, the mechanistic implication is that deactivating ROCK2 will increase endosomal SORL1-retromer.

ROCK2 activation is controlled by a handful of upstream RhoGEFs (Rho Guanine Nucleotide Exchange Factors)^31^. They act to molecularly switch the RhoA-GTPase from RhoA-GDP to RhoA-GTP, which then binds to and activates ROCK2 primarily through a conformational change that relieves ROCK2’s internal "auto-inhibitory" state^32^.

For several reasons, and with an eye towards therapeutics, we set out confirm the mechanistic prediction using an established pharmacological inhibitor that deactivates ROCK2 by blocking the ability of RhoGEF12, a dominate ROCK2 activator^33^, to bind RhoA^34^. Besides the fact that it is one of the few RhoGEFs found enriched in synapses^35^, several studies suggest that RhoGEF12 is upregulated in AD^36–38^, including a re-examination of a dataset from our previous study that first linked retromer to SORL1 by probing for molecules differentially expressed in AD’s vulnerable brain region^39^. Because of functional redundancy, inhibiting a single RhoGEF is predicted to have high safety profiles^40^, a prediction has been borne out for RhoGEF12. Deactivating ROCK2 by inhibiting or depleting RhoGEF12 in mouse models has not only already been shown to be therapeutic against various health stressors^41–44^, but it also does so safely, even when RhoGEF12 is completely knocked out. Moreover, since dozens of ROCK2 inhibitors have already been developed, if RhoGEF12 is validated as a therapeutic target, it would open up a novel upstream avenue for drug discovery and development.

By using the RhoGEF12 pharmacological inhibitor in a series of mouse and human neuronal culture studies, we validate both the predicted mechanism and RhoGEF12’s therapeutic potential. When reviewed together with other studies, the results offer mechanistic insight into how endosomal SORL1-retromer and the recycling pathway it regulates is normally controlled in the brain, and how this mechanism can be therapeutically co-opted for AD and potentially other neurodegenerative disorders.

## RESULTS

### ROCK2-dependent SORL1 phosphorylation reduces the binding affinity of SORL1-retromer

The first step was an *in vitro* analysis of synthesized peptides that cover two of SORL1’s putative ROCK2 binding sites. As illustrated in Figure 2A, one is at Ser 2206, located at the C-terminal end of the cytosolic tail, and for which some experiments suggest that this site may also be a phosphorylation site ^25^. The other is at Ser 2167, at the N-terminal end of the tail, which, based on PhosphoSite Plus (https://www.phosphosite.org/homeAction.action), is another putative ROCK2 binding and phosphorylation site.

**Figure 2.**
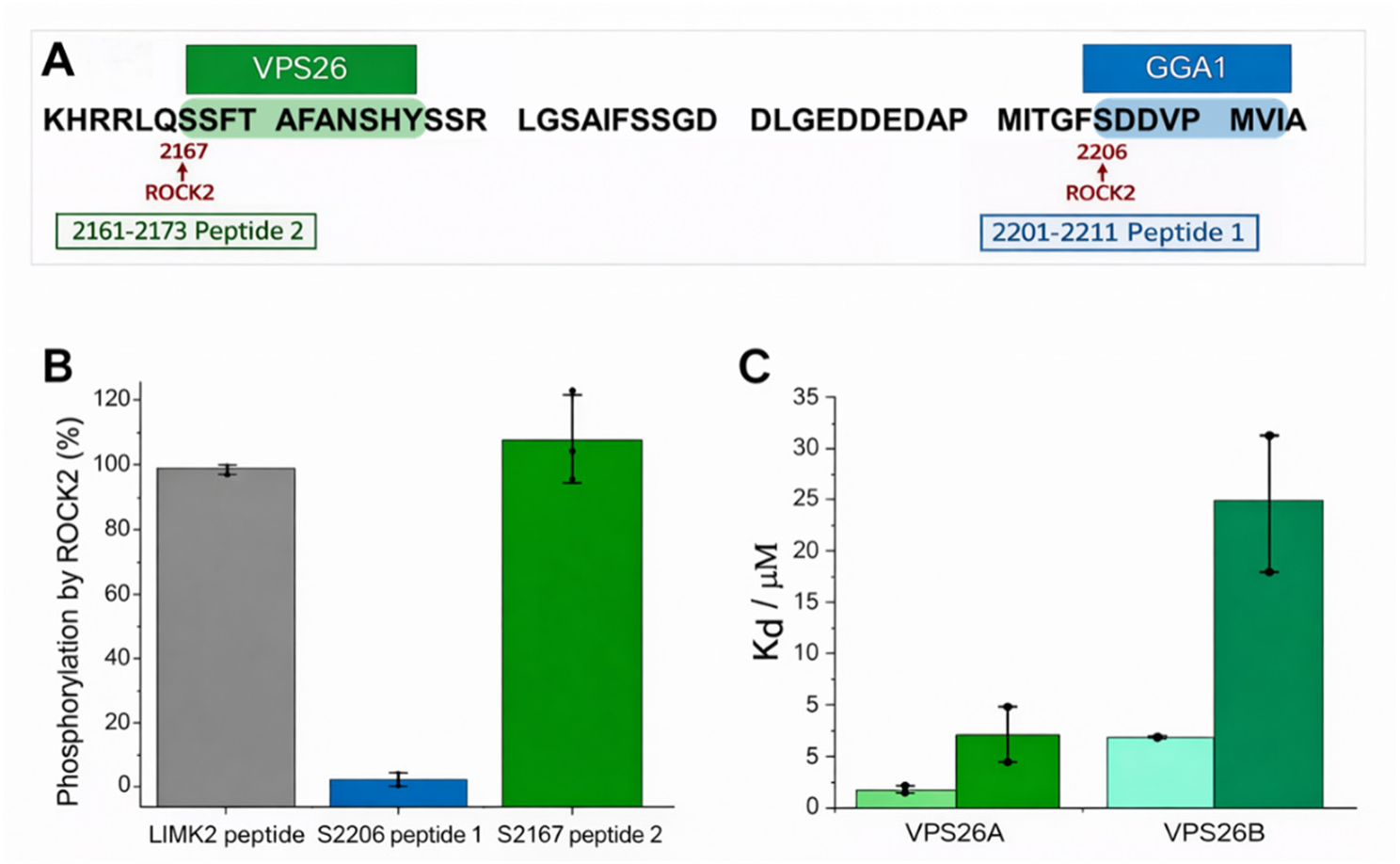
ROCK2-dependent SORL1 phosphorylation reduces the affinity of SORL1-retromer binding. **A.** SORL1’s multiple domains are illustrated with its cytoplasmic tail enlarged to highlight its GGA1, and VPS26 binding motifs. The two peptides that were investigated *in vitro* and which cover two of SORL1’s tail putative ROCK2 phosphorylation epitopes at Ser 2206 and at Ser 2167 are marked blue and green respectively. **B.** By incubating ROCK2 with the two peptides, as well as with a LIMK2 kinase peptide as a reference, ROCK2 phosphorylation at the SORL1 serine 2167 epitope shows high phosphorylation (107% of LIMK2 peptide), while serine 2206 epitope is only marginally phosphorylated (7% of LIMK2 peptide). **C.** Equilibrium binding constants between non-phosphorylated (left half-column) and phosphorylated (right half-column) SORL1 Peptide 2 (containing Ser2167 phosphorylated by ROCK2) and VPS26 variants measured by MST, shows that ROCK2-dependent phosphorylation of S2167 reduces the SORL1 peptide’s affinity to VPS26A/26B by 4-5 fold.

By incubating ROCK2 with both of these peptides and also with the LIM2 kinase canonical peptide that is a high-affinity potent substrate for ROCK2^45^, we find that both sites are substrates but there is much higher phosphorylation at the tail’s N-terminal S2167 site (compared to the 2206 site), equivalent to or even higher than the phosphorylation of the LIM2 kinase peptide (Fig. 2B). It was verified by mass spectrometry that ROCK2 phosphorylates SORL1 at Ser2167. Notably, the 13-residue peptide containing the S2167 binding site to ROCK2 overlaps with the known binding site for VPS26 (Fig. 2A). VPS26 has two paralogues, VPS26A and VPS26B, and so we performed microscale thermophoresis (MST) with the unphosphorylated and phosphorylated (pS2167) peptides and purified ROCK2, VPS26A, and VPS26B. Compared to unphosphorylated S2167 peptide, for which the ROCK2 Kd=9.0±4.6 µM, the binding affinity for ROCK2 increased when S2167 is phosphorylated (Kd=3.8±1.8 µM). Most importantly, upon phosphorylation, the affinity for the SORL1 S2167 peptide decreases approximately 4-5 fold (Fig 2C) for both VPS26B (Kd=6.4±0.1 µM for unphosphorylated and 26±6 µM for phosphorylated S2167 peptide) and VPS26A (K_d_=1.4±0.3 µM for unphosphorylated and 6.3±2.3 µM for phosphorylated S2167 peptide).

Together, the *in vitro* studies uncovered a novel ROCK2 phosphorylation site on SORL1’s cytoplasmic tail, that when phosphorylated decreases SORL1’s binding affinity to both VPS26A and VPS26B. When this site on SORL1’s tail is phosphorylated by ROCK2, the diminished SORL1 binding for VPS26B is more pronounced. With relevance to AD, previous studies have shown that, compared to VPS26A, VPS26B differentially controls Aβ40 and Aβ42 production and is linked to the recycling pathway^46^.

### RhoGEF12 Inhibition increases endosomal SORL1-retromer

It was important to test the mechanistic prediction that inhibiting RhoGEF12 will increase endosomal SORL1-retromer in primary wildtype neurons by relying on endogenous proteins, since expressing exogenous constructs can alter intracellular protein localization. Despite the fact that in wildtype neurons SORL1 and retromer are found to partially co-localize to endosomes^27–29^, we used confocal immunocytochemistry to test whether the selective RhoGEF12 inhibitor Y16^34^ increases endosomal SORL1-retromer. We first administered 3uM and 5uM of Y16 to primary mouse cultured neurons for 72 hours and by immunoblotting find that, compared to DSMO control, SORL1 levels were slightly increased with 5uM Y16 treatment (Supplemental Fig. 1A). Then, we replicated two confocal immunocytochemistry experiments, using SORL1 antibodies, VPS35 antibodies as an indicator of retromer, and EEA1 antibodies as an indicator of endosomes Compared to DMSO control, 5uM of Y16 applied for 72 hours resulted in a modest but reliable increase in the colocalization of VPS35 with EEA1; VPS35 with SORL1; and SORL1 with EEA1 (Fig. 3).

**Figure 3.**
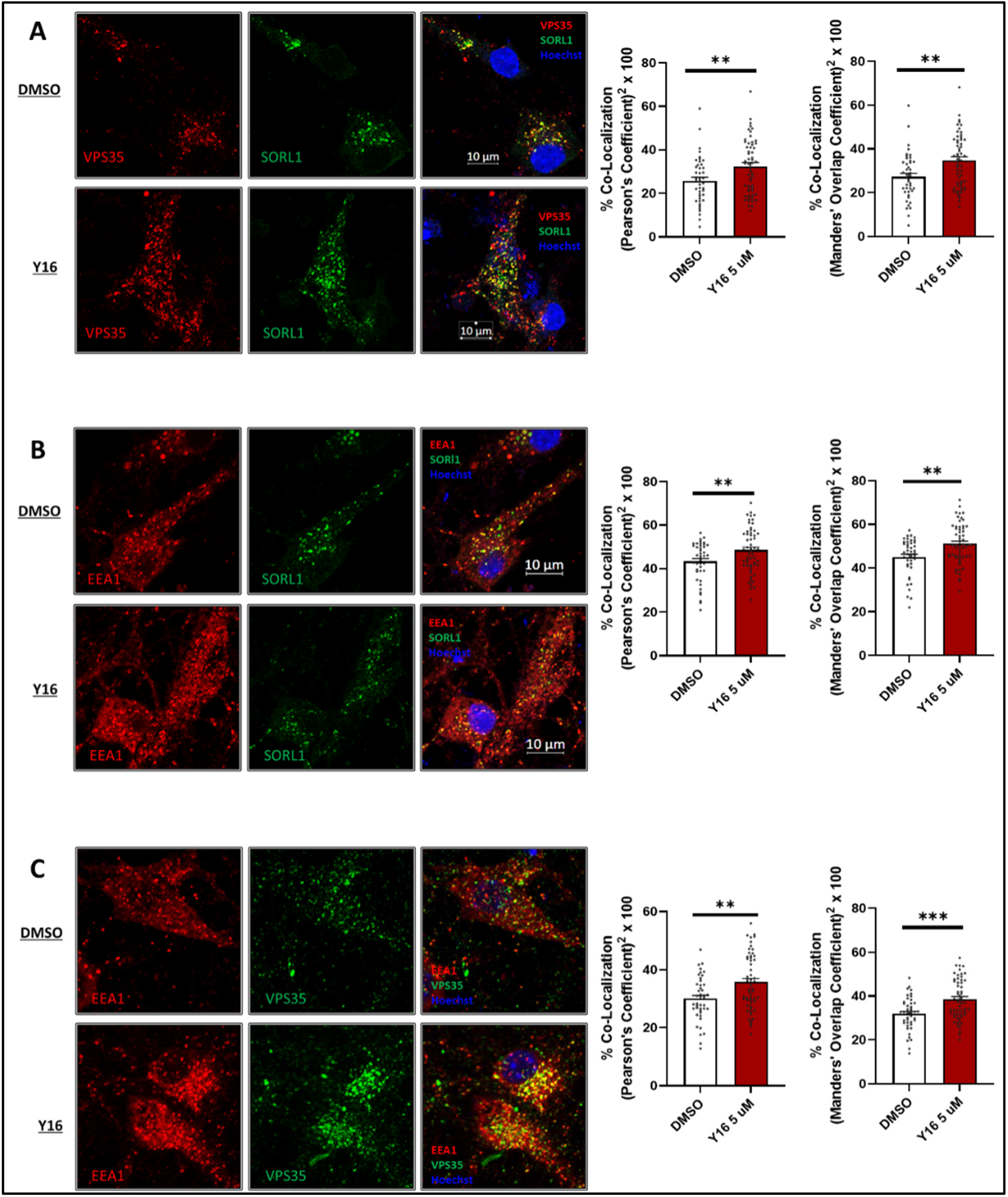
RhoGEF12 inhibition increases the colocalization of endosomal SORL1-retromer. Compared to DMSO control, applying the RhoGEF12 pharmacological inhibitor Y16 to mouse cultured neurons increased the colocalization of VPS35 with SORL1 **(A)**, SORL1 with the endosomal marker EEA1 **(B)**, and VPS35 with EAA1. as quantified by a Pearson’s coefficient (left bar graphs) and a Mander’s Overlap coefficient (right bar graphs. ** indicates p<0.01; *** indicates p<0.001).

To further confirm the conclusion that Y16 increases endosomal SORL1-retromer, we optimized a fractionation protocol that can assay cytosolic vs. membrane-bound proteins (Fig. 4A) and validate the protocol using proteins that are known to reside primarily in the cytosol or on membranes, including endosomal membranes (Supplemental Fig. 2). By immunoblotting we find that, compared to DMSO control, 5uM of Y16 applied for 72 hours increased VPS35’s distribution to the membrane fraction (Fig. 4B).

**Figure 4.**
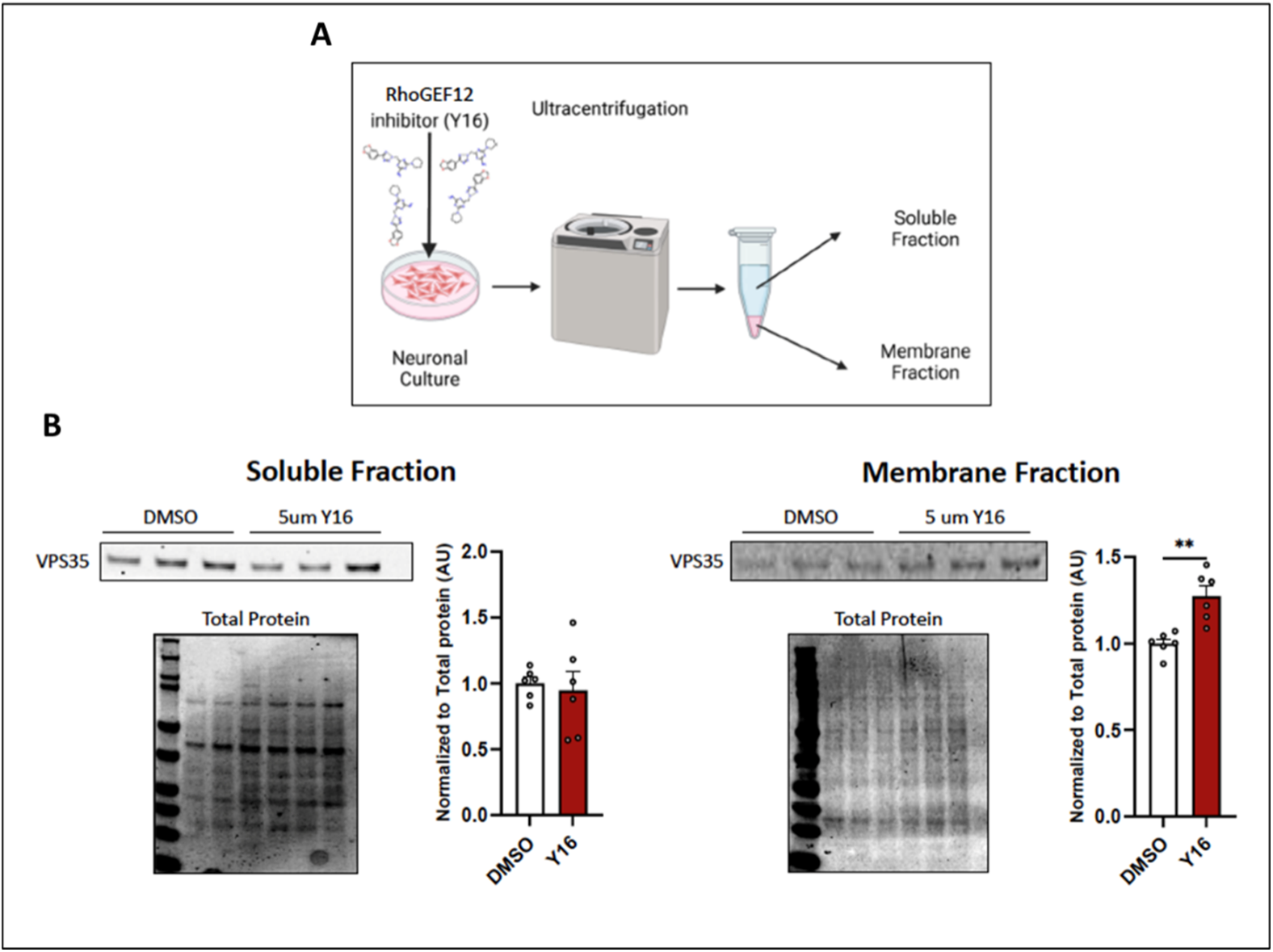
RhoGEF12 inhibition increases retromer’s membrane fraction. **A.** An illustration of the soluble versus membrane fractionation experimental protocol. **B.** Compared to DMSO control, applying the selective RhoGEF12 pharmacological inhibitor Y16 to mouse cultured neurons results in a reliable shift of retromer (as indicated by VPS35) from the soluble fraction to the membrane fraction (* indicates p <0.05, ** indicates p<0.01).

### RhoGEF12 Inhibition reduces Aβ secretion in a SORL1-dependent manner

SORL1-retromer endosomal recycling controls the production and secretion of Aβ40 and Aβ42 by recycling APP out of endosomes^11^, and so a joint reduction in both Aβ40 and Aβ42 has emerged as one of the most reliable indicators of pathway upregulation, including in human iPSC-derived neurons^8,22,47^. As illustrated in Figure 5A, we therefore generated a collection of human iPSC-derived neurons (iNeurons) and completed two experiments.

**Figure 5.**
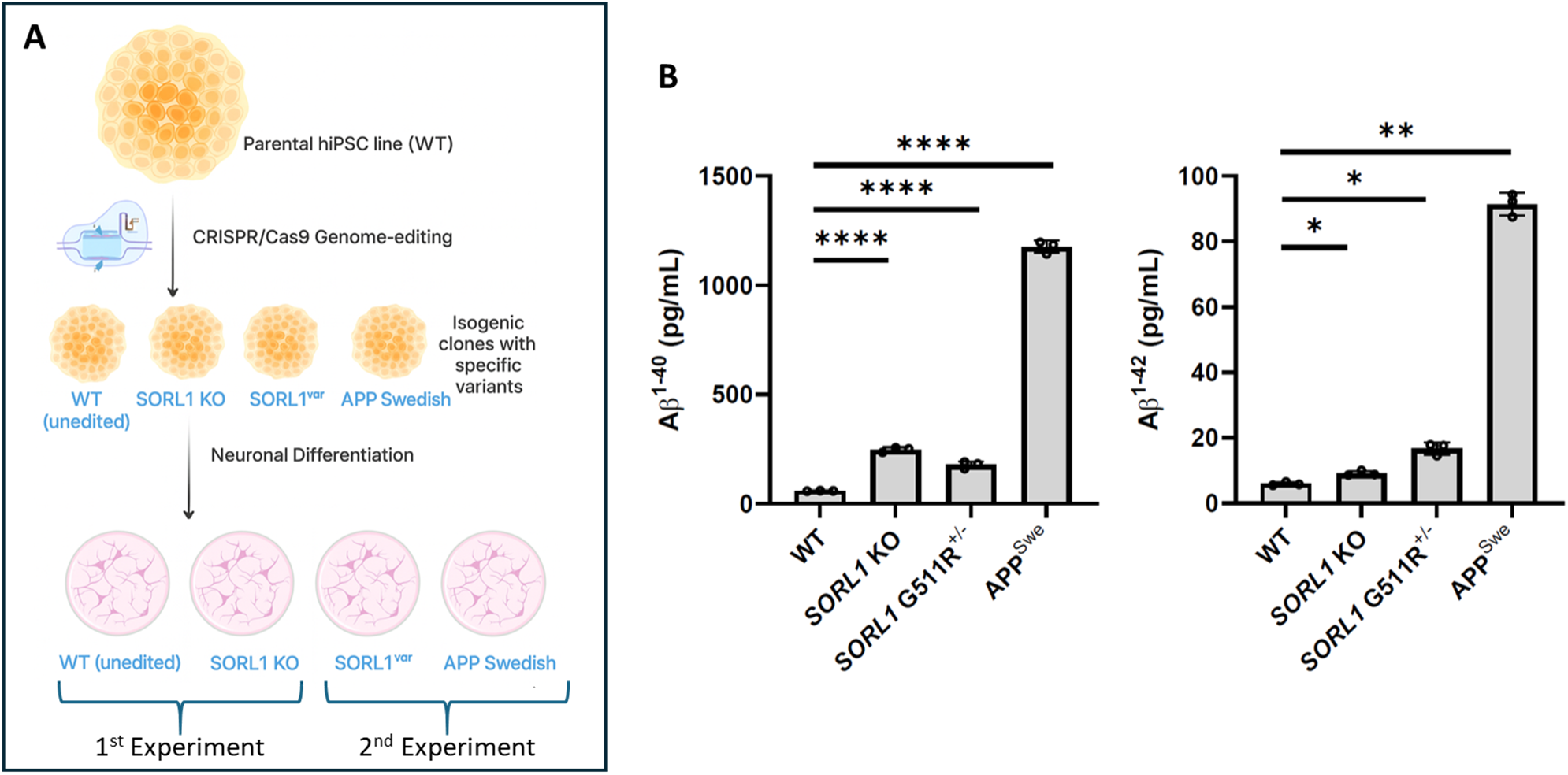
Human iPSC-derived neuronal lines. **A.** An illustration of how the four human iPSC-derived neurons (iNeurons) were generated. **B.** Confirming previous observations, compared to ‘wildtype’ controls, each line shows an increase in Aβ40 (left bar graph) and Aβ42 (right bar graph). (* indicates p<.05, ** indicates p<0.01; **** indicates p<0.0001).

The first experiment was designed to confirm the mechanistic conclusion that RhoGEF12 inhibition upregulates the pathway in a SORL1-dependent manner. To test this prediction, we generated SORL1 knockout (SORL1^⁻/⁻^) iNeurons, previously shown to have increased Aβ40 and Aβ42 production and secretion^22^ and their isogenic ‘wildtype’ parental iNeurons. We confirm that, compared to the parental wildtype iNeurons, SORL1 depleted iNeurons show an elevation in Aβ40 and Aβ42 assayed from the cultured media (Fig. 5B&C). We additionally confirm the findings in mouse cultured neurons, showing that in wildtype iNeurons, Y16 treatment results in a small increase in SORL1 (Supplemental Fig. 1B).

We then administered Y16 for 72 hours at three ascending doses: 5uM, 10uM, and 30uM. ELISA was used to assay cultured media for Aβ40 and Aβ42. No evidence of toxicity was observed at any dose. A multivariate ANOVA was used in which Y16 dose was included as the fixed factor, with Aβ40 and Aβ42 included as the dependent variables. In the ‘wildtype’ iNeurons, a reliable general effect was observed with Y16 reducing Aβ40 (F=4.47, p=0.032) and Aβ42 (F=5.76, p=0.015). By applying a simple contrast, compared to 5uM Y16, 10uM Y16 was observed to reliably reduce Aβ40 (p=0.019) and Aβ42 (p=0.011), and 30uM Y16 was observed to reliably reduce Aβ40 (p=0.027) and Aβ42 (p=0.012). While no reliable differences were observed when comparing 10uM to 30uM Y16, a polynomial contrast revealed a linear reduction in Aβ40 (p=0.027) and in Aβ42 (p=0.012), suggesting a dose-dependent effect (Fig. 6A). When the same ANOVA model was applied to SORL1 depleted iNeurons, no reliable general effect was observed for Y16 on Aβ40 (F=0,81, p=0.46) or on Aβ42 (F=0.7, p=0.51), and all contrasts were null (Fig. 6B).

**Figure 6.**
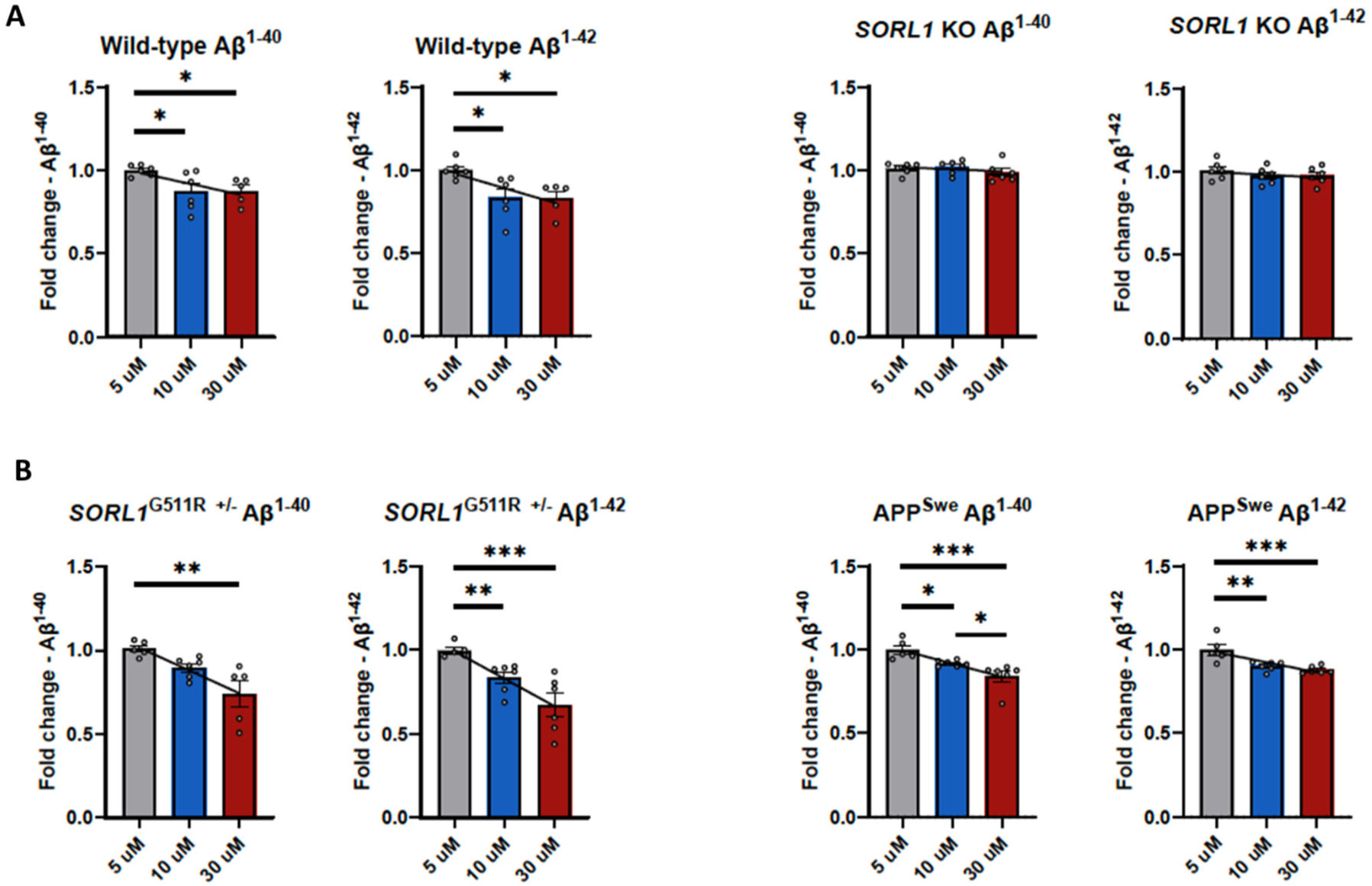
RhoGEF12 Inhibition reduces Aβ40 and Aβ42 secretion. **A.** In ‘wildtype’ human iPSC-derived neurons (iNeurons) treatment with the RhoGEF12 pharmacological inhibitor Y16 shows a dose-dependent reduction in Aβ40 and Aβ42 (left panels), but no effect in SORL1 depleted iNeurons (right panels). **B.** Treatment with the RhoGEF12 pharmacological inhibitor Y16 shows a dose-dependent reduction of Aβ40 and Aβ42 in iNeurons expressing an AD-associated SORL1 mutation (left panels) and in iNeurons expressing an AD-associated APP mutation (right panels). p < 0.05*, p < 0.01**, p < 0.001***, p < 0.0001****; lines represent significant regression relationships across doses.

Collectively, these results support the mechanistic conclusion, showing that RhoGEF12 upregulates SORL1-retromer endosomal recycling in a SORL1-dependent manner.

### RhoGEF12 Inhibition reduces Aβ secretion in AD human neuronal models

The second iNeuron experiment was designed to test the therapeutic potential of RhoGEF12 inhibition. As illustrated in Figure 5A, to model patients carrying SORL1 pathogenic mutations, we used CRISPR-engineered neurons expressing the pathogenic SORL1^G511R^ variant, which although it delivers SORL1 to endosomes, results in increased Aβ40 and Aβ42 production^21^ and secretion as we confirm (Fig. 5B). To model patients carrying APP pathogenic mutations we used CRISPR-engineered neurons expressing the Swedish APP mutation (APP^SWE^), which results in increased endosomal Aβ40 and Aβ42 production and secretion^22^, which we also confirm (Fig. 5A&B).

Y16 was administered for 72 hours using the same ascending doses and ELISA was used to assay cultured media for Aβ40 and Aβ42. No evidence of toxicity was observed at any dose. The same multivariate ANOVA model was used, in which Y16 dose was included as the fixed factor and with Aβ40 and Aβ42 included as the dependent variables. In the SORL1 mutant iNeurons, a reliable general effect was observed with Y16 reducing Aβ40 (F=8., p=0.005) and Aβ42 (F=10.2, p=0.002). By applying a simple contrast, compared to 5uM Y16, 10uM Y16 did not reliably reduce Aβ40 (p=0.11) but did reliably reduce Aβ42 (p=0.010), and 30uM Y16 was observed to reliably reduce Aβ40 (p=0.002) and Aβ42 (p<0.001). Compared to10uM, 30uM was observed to reliably reduce Aβ40 (p=0.03) and to result in a trending reduction in Aβ42 (p=0.069). A polynomial contrast confirmed a dose-dependent effect with a linear reduction in Aβ40 (p=0.002) and in Aβ42 (p<0.001) by dose (Fig. 6C).

In the APP mutant iNeurons, a reliable general effect was observed with Y16 reducing Aβ40 (F=9.9, p=0.002) and Aβ42 (F=10.8, p=0.001). By applying a simple contrast, compared to 5uM Y16, 10uM Y16 reliably reduced Aβ40 (p=0.039) and reliably reduced Aβ42 (p=0.003), and 30uM Y16 was observed to reliably reduce Aβ40 (p<0.001) and Aβ42 (p<0.001). Compared to 10uM, 30uM was observed to reliably reduce Aβ40 (p=0.039) with no effect on reducing Aβ42 (p=0.39). Nevertheless, a polynomial contrast suggested a strong dose-dependent effect with a linear reduction in Aβ40 (p<0.001) and in Aβ42 (p<0.001) (Fig. 6D).

Collectively, the results of this iNeuron experiment provide preclinical validation that inhibiting RhoGEF12 is a therapeutic target in AD.

### SORL1 deficiency elevates ROCK2 in the mouse brain

A genetic association study first found that, among retromer-related proteins, SORL1 is the one with the strongest genetic linkage to late-onset AD^48^, and subsequent large-scale GWAS studies have established that SORL1 is one of the most common genes implicated in late-onset AD^49^. More recent genetic studies have established that SORL1 deficiency^50^ and loss of endosomal SORL1^51^ are causally pathogenic, while SORL1 overexpression appears protective in late-onset AD^52^. Concordantly, numerous postmortem studies have reported that SORL1 is deficient in the brains of late-onset AD patients^46,53^. Independent studies have also indicated that ROCK2^54^ levels are elevated in sporadic AD.

To investigate whether SORL1 reductions in the brain might be linked to ROCK2 elevations, we probed the cortex of mice expressing a targeted gene deletion of the 5’ region of *SORL1*’s exon4, resulting in SORL1 deficiency^53^. By immunoblotting, we found that, compared to wildtype littermates, ROCK2 is elevated in the SORL1 deficient mice (p<0.01) (Fig. 7).

**Figure 7.**
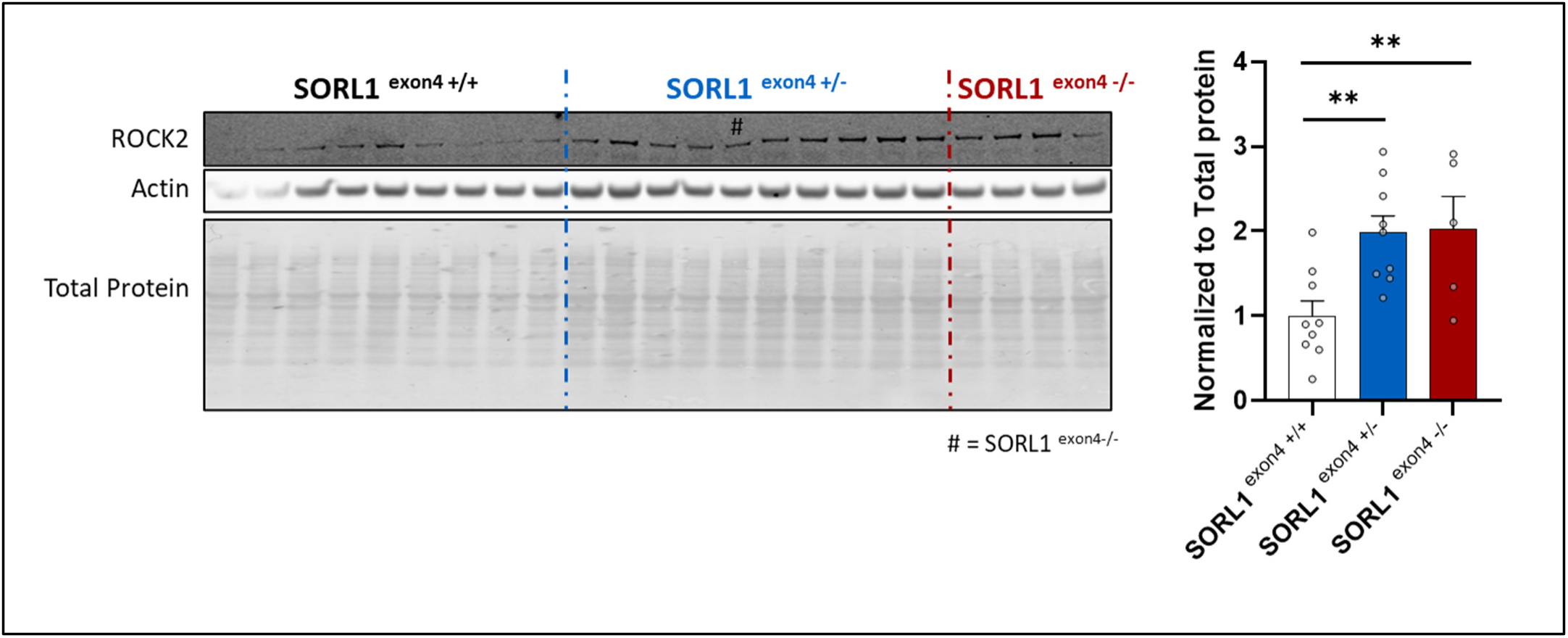
SORL1 deficiency leads to an elevation of ROCK2 in the mouse brain. Compared to wildtype littermates (SORL1^exon4+/+^), SORL1 deficient mice (SORL1^exon4-/+^ or SORL1^exon4-/-^) show an approximate doubling in ROCK2 levels in the brain. ** indicates p<0.01.

While these findings suggest that SORL1 deficiency can drive the ROCK2 elevations observed in AD brains, the precise mechanism driving this association remains unknown. By whichever mechanism, based on our findings above, the ROCK2 elevation triggered by SORL1 deficiency will further accentuate a reduction in endosomal SORL1-retromer, setting up a potential vicious cycle.

## DISCUSSION

Our findings have both mechanistic and therapeutic implications. Mechanistically, our detailed biochemical analysis of SORL1’s cytoplasmic tail established that it has multiple ROCK2 phosphorylation sites, and that one site’s high level of phosphorylation suggests that the affinity with which ROCK2 interacts with SORL1 is higher than most other known ROCK2-interacting proteins. This conclusion agrees with a study that performed a screen on proteins pulled down by SORL1, finding that, besides itself, ROCK2 was SORL1’s top binding partner^25^. Ultimately, it is the functional consequences of ROCK2-dependent SORL1 phosphorylation that are most important. Our results show how this phosphorylation can control SORL1’s retromer binding by reducing the affinity of SORL1 for VPS26.

Collectively, our results uncover a novel mechanism establishing how endosomal SORL1-retromer can be actively regulated. Moreover, the fact that both ROCK2 and SORL1 are enriched in the brain, taken together with their distinctly strong and functionally impactful interaction, allows the conclusion that ROCK2-dependent SORL1 phosphorylation can act as a dominant regulator of the brain’s SORL1-retromer endosomal recycling pathway.

We co-opted this mechanistic insight into our therapeutic goal for developing SORL1-based interventions and provide preclinical evidence that RhoGEF12 is a therapeutic target. Future studies can cross RhoGEF12 knockout mice, which have been generated and found to be benign^41^, with mice modeling neurodegenerative disorders to characterize the full dynamic range of AD-relevant readouts. These mice studies will be useful to better determine the effect sizes Aβ reduction when RhoGEF12 is targeted *in vivo* and for longer durations. Beyond accelerated Aβ production, mouse models with retromer deficiency are beginning to show evidence of neurodegeneration^55^, and so future studies can test whether RhoGEF depletion ameliorates the neurodegenerative process. It is neuronal loss, after all, that defines neurodegenerative disorders, and which results in the devastating cognitive morbidity of patients.

Additionally, future studies in model systems can test whether the therapeutic benefits of targeting RhoGEF12 extend beyond AD. A recent study^16^ showed that tau aggregates sequester retromer proteins, effectively ‘stealing’ them away from endosomal trafficking. This tau aggregate sequestration was found not just in AD but in other primary tauopathies such as FTD, corticobasal degeneration (CBD), and progressive supranuclear palsy (PSP)^16^. Another recent study showed that cytoplasmic TDP-43 aggregates can result in retromer deficiency^17^. Besides commonly occurring as a co-pathology in AD^56^ and in approximately 50% of FTD, cytoplasmic TDP-43 aggregates are found in nearly all patients with amyotrophic lateral sclerosis (ALS)^57^. Indeed, previous studies have observed retromer deficiency in TDP-43 positive ALS spinal neurons of patients^18^ and in TDP-43 positive ALS iNeurons^19^.

Most importantly for therapeutics, by providing evidence that RhoGEF12 is a therapeutic target, our findings justify future medicinal discovery and development. The previous study that identified the selective RhoGEF12 pharmacological inhibitor used here relied on known crystal structures of RhoGEF12 interacting with RhoA to screen for small molecule protein-protein inhibitors^58^. Thus, it is possible to screen and develop additional small molecules that might have better medicinal properties. Moreover, antisense oligonucleotides (ASOs), small interfering (siRNAs) or small hairpin RNAs (shRNAs) directed at reducing brain RhoGEF12 are alternative strategies for drug discovery.

In conclusion, mechanistically and preclinically validating RhoGEF12 as a therapeutic target will hopefully inspire future drug discovery, so that they might ultimately become human drugs for a range of common and still undertreated neurodegenerative disorders.

## MATERIALS AND METHODS

### In Vitro Studies

#### Protein expression and purification

The gene construct for expressing the truncated (residues 1-420) version of human ROCK2 kinase is based on synthetic genes produced at Geneart (Regensburg, Germany) cloned into the pONE30A vector^59^ between NheI and NotI cleavage sites. The DNA sequence of the synthetic gene is based on Uniprot entry O75116. The genes were codon optimized for expression in baculovirus-insect cells. The ROCK2 protein (containing MBP tags at the N-terminus) was expressed^32^ in Sf9 insect cells after co-transfection using flashBAC GOLD baculoviral expression system (Oxford Expression Technologies, Oxford, UK). After three rounds of virus amplification, insect cells at 2 × 10^6^ cell/mL density were infected with the recombinant baculovirus to express the protein in Insect-XPress medium (Lonza, Basel, Switzerland) for 3 days at 27 °C in a shaker flask at 220 rpm. The cells were centrifuged and resuspended in 25 mM HEPES, 500 mM NaCl, pH 7.4 buffer containing cOmplete ULTRA (Roche, Basel, Switzerland) protease inhibitor cocktail mix and 1 mM DTT, then sonicated. Cell debris was centrifuged and discarded, and the proteins were purified from the supernatant using amylose affinity chromatography. The bound proteins were eluted by the elution buffer (binding buffer plus 10 mM maltose). During affinity chromatography, peak fractions were collected and analyzed with SDS-PAGE and Western blot. The collected protein fractions were subjected to gel-filtration using Sepharose G25 to 10 mM Hepes, 150 mM NaCl buffer with the pH of 7.4 and were concentrated to approx. 2 mg/ml in Amicon Ultracel (Merck Group, Darmstadt, Germany) – 10K centrifugal filter cells.

#### Peptide synthesis

All peptides were synthesized by Genscript (Piscataway, NJ, USA) and the purities were ≥95%. Peptide1 (2201-2211), and peptide 2 (2161-2173) represent the 2201-2211 (MITGFSDDVPM) and 2161-2173 (KHRRLQSSFTAFA) residues of SORL1 (based on Uniprot entry Q92673 numbering). Peptide 2P has the same sequence as peptide 2 but has a synthetical phosphoserine at position 7 (representing residue 2167 of the original SORL1 sequence). LIMK peptide has the sequence of RKKRYTVVGNP, representing residues 500-510 of human LIMK2 (based on Uniprot entry P53671 numbering).

#### Phosphorylation studies

Phosphorylation rate was measured using ADP-Glo Kinase Assay^60^ kit (Promega Corporation, Madison, WI, USA). The reaction mixes contained 160 nM ROCK2 kinase domain, 100 µM ATP and 150 µM peptide in a reaction buffer of 50 mM HEPES (pH 7.5), 10 mM MgCl2, 1 mM EGTA, 0.01% BRIJ-35 buffer. 20 µl of the reaction mixes were incubated at 30°C for 60 minutes and 5 µl aliquots were taken to ADP-Glo measurement according to manufacturer’s protocol in a white 384-well plate. Luminescence was measured using a BMG Labtech ClarioStar PLUS (BMG LABTECH, Ortenberg, Germany) multimode microplate reader. Three independent measurements were conducted and averaged.

#### Microscale Thermophoresis

Proteins (ROCK2 kinase domain, VPS26A full-length*, VPS26B full-length*) were diluted to 10 µM prior to labeling, and gel filtrated into labeling buffer (130 mM NaHCO_3_, 50 mM NaCl at pH 8.4). Proteins labeled using Monolith Protein Labeling Kit RED-NHS 2nd Generation Red-NHS (2^nd^ generation) kit (NanoTemper Technologies GmbH, Munich, Germany) according to manufacturer’s protocol. Peptides 2 and 2P were diluted 2-fold in 20 mM Tris (pH 8.0), 100 mM NaCl, 2 mM DTT and 0.05% Triton X-100 buffer in a concentration range of 1-500 µM. The thermophoresis of each protein-peptide mix was measured at a constant protein concentration of 50 nM using 20% LED energy and 40% MST energy in the Monolith NT.115 instrument at red fluorescence channel at 25°C. The T-Jump phases of the thermophoresis curve were applied for affinity calculation. K_d_ values were calculated with MO.affinity, the instrument’s native software^61^.

### Primary Mouse Neuronal culture

Postnatal day 0 primary mouse cortical and hippocampal neuronal cultures from C57BL/6J mice were implemented as described previously ^7,62^. Cultures were kept at 37 °C in 5% CO_2_ incubator. Neurons (1.2E+06 per well) were allowed to mature for 14 days after plating in a 6-well plate. At day 14 the cells were treated with 5uM Y16; DMSO treatment was used as control. The drug was replenished at day 16. The culture was maintained for 3 days (72 hours) after initial treatment and harvested at day 17. Cultured primary neurons for ICC experiments were plated and treated in the same manner with the following exceptions: 1- Cells were plated on polyornithine coated coverslips. 2- 24 well plate was used instead of 6 well plate. The neurons were fixed using 4% PFA and stored for staining.

#### Membrane isolation

Only freshly harvested samples were used and processed at 4 °C unless otherwise noted. An assay buffer (AB) containing 50 mM HEPES pH 8.0, 150 mM NaCl, 2 mM MgCl₂, HALT protease inhibitor (1×), was prepared for homogenization. Tissue/cell material was homogenized in AB + phosSTOP using motorized blue pestles in Beckman Coulter ultracentrifuge tubes (Ref. 357448) to ensure proper rotor sealing. Homogenates were briefly checked at 90,000 × g for 3 min, then subjected to ultracentrifugation at 100,000 × g for 25 min (4 °C). The supernatant (soluble fraction) was collected carefully to avoid particulate carryover. The pellet (membrane fraction) was resuspended in AB containing DNase (prepared at 40 µL DNase per 2 mL AB, scaled to match the homogenization volume) and incubated at room temperature for 30 min to remove nucleic acids.

Following DNase treatment, membrane suspensions were clarified by a second ultracentrifugation at 100,000 × g for 25 min and the supernatant discarded. Pellets were rinsed three times by resuspension in AB and repeat centrifugation (100,000 × g, 15 min) to remove residual soluble material. Membrane pellets were then solubilized in RIPA buffer supplemented with phosSTOP and roche cOmplete mini protease inhibitor and homogenized again; these extracts were centrifuged at ∼16,100 × g for 15 min and the resulting supernatant retained as the final membrane fraction. Protein concentrations for all soluble and membrane fractions were determined by BCA assay and samples prepared for SDS–PAGE and downstream analyses.

#### Western blots

Proteins from neuronal cultures and the mouse brain were isolated as described previously^7^. Lysates from the samples were run on NuPAGE® Bis-Tris 4-12% gels, washed in 20% ethanol for 10 minute, transferred onto nitrocellulose membranes using iblot and were probed with antibodies. Primary antibodies targeting the following proteins were used for probing: VPS35 (ab57632, Abcam, 1:1k), β-actin (ab6276, Abcam), Tau (A0024, Dako 1:10k) GluA1 (mab2263, Millipore, 1:1k), APP-CTF (ab32136, Abcam, 1:10K & 1:2k respectively), ROCK2 (ab125025, Abcam, 1:5k). The primary antibodies were selected based on extensive citations by previous studies; most of the antibodies used are monoclonal antibodies with minimal non-specific signal and several of them have been validated by KO testing. IRDye® 800 or 680 antibodies (LI-COR) were used as secondary with dilutions of 1:10k for 800CW, 1:15k for 680RD, and 1:25k for 680LT antibodies. Western blots were scanned using the Odyssey imaging system as described previously ^63^.

#### Fluorescent Immunocytochemistry (ICC)

Standard staining and microscopy techniques were used with some modifications. All washes were done with PBS (1X-PBS-triton-x containing 0.01% Sodium Azide). Briefly, antigen retrieval was performed using Citric Acid buffer (10mM sodium citrate, 1.9mM citric acid, pH6.0). Since these are cells, the antigen retrieval step has to be very gentle. Buffer was preheated in a covered 50ml beaker in a water bath to 98°C before immersing the coverslips for 10 minutes. The coverslips were monitored to keep the buffer from boiling. Buffer and coverslips were allowed to cool down at room temperature for 30min. After one wash coverslips were permeabilized using 0.01% digitonin (wt/vol) for 10 min and then blocked in 5% donkey serum in PBS-triton-x (vol/vol) for 1.5-3 hours, followed by incubation in primary antibodies overnight at 4°C, and for an additional 1 hour at room temperature the next morning. Primary antibodies were prepared in 1% donkey serum (in PBS-triton-x) for the following proteins: VPS35 (ab10099, abcam, 1:500), Sorl1 (anti-LR11 BD-611861, BD bioscience 1:1k). Next day after five washes, the coverslips were incubated overnight for SORL1 with secondary antibody at room temperature. Secondary antibodies (ThermoFisher) conjugated with alexa fluor dyes were used at a dilution of 1:250 in 1% donkey serum (in PBS-triton-x). After 3 washes Hoechst 33258 nuclear stain (ThermoFisher H3569, 1:1500 for 5 min) was used to counter stain the nuclei. After another five washes coverslips were mounted onto the slides with ProLong Gold Antifade Mountant (ThermoFisher P36934).

### Human iPSC-derived neurons

The generation of the hiPSC cell lines used in this study has been previously described^21,22,64^. In brief, SORL1 KO, heterozygous SORL1 G511R, and homozygous APP Swedish status were generated utilizing CRISPR/Cas9 gene editing in the CV background cell line. This cell line is male and has the APOE E3/E4 genotype. All cell lines used in this study have normal karyotype and were routinely rested for mycoplasma (MycoAlert).

For the SORL1 series and APP Swedish mutation, hiPSCs were differentiated into neurons as previously described^21,29,64^. hiPSCs were differentiated into cortical neurons using dual SMAD inhibition. hiPSCs were maintained on Matrigel (Corning cat. #35631) coated culture plates in mTesR Plus (StemCell cat #100-0276) medium. On day 1 of neural induction, cells were exposed to dual SMAD inhibitors 10μM SB-431542 (Biogems cat. #BG6675SKU301) and 0.5μM LDN-193189 (Biogems cat. #BG5537SKU106) in basal neural maintenance medium (BNMM): 1:1 DMEM/F12 + glutamine (Gibco cat. #11320-033) and Neurobasal (Gibco cat. #21103-049), supplemented with 1% B27 (Gibco cat. #17504-044), 0.5% N2 (Gibco cat. #17502-048), 0.5% GlutaMax (Gibco cat. #35050-061), 0.5% insulin-transferrin-selenium-A (Gibco cat. #51300-044), 0.5% Non-essential amino acids (NEAA) (Thermo Fisher cat. #11140050), 0.2% β-mercaptoethanol (Thermo Fisher cat. #21985023). Medium was fully exchanged and fed daily for seven days. On day eight, cultures were dissociated with Versene (Gibco cat. #15640066) and passaged to fresh Matrigel-coated culture plates. The next day, cells were fed with BNMM + 20 ng/ml fibroblast growth factor (FGF) (R&D Systems cat. #233-FB/CF). On day sixteen, cells were passaged into fresh Matrigel-coated culture plates. Cells were maintained with BNMM + FGF until day twenty-three, where they were sorted using fluorescence-activated cell sorting (FACS) flow cytometry to isolate neural progenitor cells (NPCs). NPCs were identified with a quad cell surface immunostain: positive for CD24 and CD184 while negative for CD44 and CD 271^65^. Following NPC isolation, they were cryopreserved or expanded for neuronal differentiation. For neuronal differentiation, eight million NPCs were plated on Matrigel-coated 10cm culture dishes and medium was changed to BNMM supplemented with 0.2 μg/ml brain-derived neurotrophic factor (BDNF) (PeproTech cat. #450-10), 0.2 μg/ml glial-derived neurotrophic factor (GDNF) (Peprotech cat. #450-02), and 500 μM db-cAMP (Sigma-Aldrich #D0627). Medium was fully replaced twice a week for three weeks. After three weeks of neuronal differentiation, cells were dissociated with Accutase (Innovative Cell Technologies cat. #AT-104). Cell suspension concentrations and viabilities were counted using Trypan blue (Gibco cat. #15250-061) and a TC20 cell counter (Bio-Rad). Cells were then replated according to experimental needs. All cells were maintained at 37° C and 5% CO2.

To assay secreted Aβ protein, differentiated neurons were plated on Matrigel-coated 96well plates at a density of 250,000 cells per well. 72 hours after passaging, neurons were treated with Y16 compound or DMSO vehicle control. All treatment and control conditions were performed in triplicate using two independent hiPSC neuron differentiations. On the day of Y16 treatment, neuronal medium was fully aspirated and replaced with 150 µl of BNMM supplemented with 0.2 μg/ml BDNF, 0.2 μg/ml GDNF, 500 μM db-cAMP, and 5, 10, or 30 µM Y16 compound. All concentrations contained equal dilutions of DMSO. Following treatment, neurons were cultured for 72 hours before harvesting. Neuronal health was monitored daily.

72-hour conditioned medium was collected for Aβ assay. Secreted Aβ levels were measured from conditioned medium using an MSD Aβ V-PLEX ELISA kit (Meso Scale Discovery cat K15200E-2). ELISA kits were performed and concentrations measured per manufacturer’s instructions. All samples were tested with two technical replicates.

### SORL1 deficient mice

We investigated mice genetically-engineered to express a targeted deletion of the 5’ region of *SORL1*. Specifically, these mice have an in-frame deletion of exon 4 of the *SORL1* gene (encoding a part of the VPS10p-domain)^66^. Previous studies have established that these mice express low levels of exon4-deleted SORL1^53^. While in these mice it is difficult to assay and discriminate the expression of exon4-deleted SORL1 from non-deleted SORL1, an examination of the brains of both SORL1^Exon4+/-^ and SORL1^Exon4-/-^ mice showed that the deletion results in a gene-dose dependent reduction in SORL1 levels and function^53^.

#### Perfusion and tissue processing

After the survival period, the mice were perfused with saline, the brains extracted, dissected, and homogenized for biochemical analysis as described above and previously^7^.

## AKNOWLEDGMENTS

## Contributions

YHQ, SK, and AG designed, strategized, and performed the experiments in mouse cultured neurons and SORL1 mice; JEY and CAW designed, strategized, and performed the experiments in human iPSC-derived neurons; GAP edited the manuscript and contributed reagents to the *in vitro* experiments, and in their interpretation and in figures generation; PZ, IH, and BV performed the *in vitro* SORL1 experiments; SAS, YHQ, JEY, and CAW analyzed the data; and SAS conceptualized and interpreted the experiments and wrote the manuscript.

