## Supplemental File for "RhoGEF12 regulates endosomal SORL1-retromer and its inhibition is therapeutic in human neuronal models of Alzheimer’s disease"

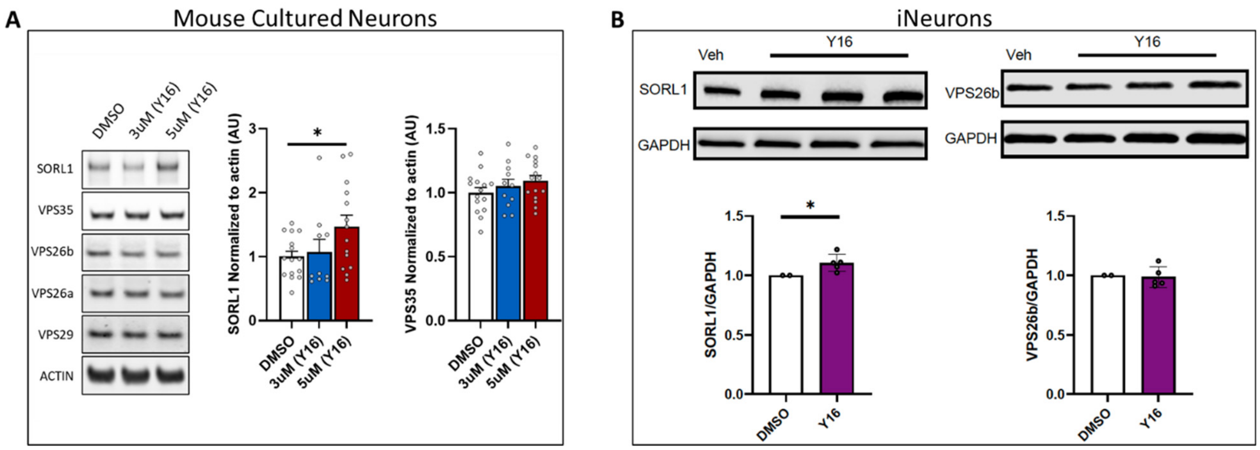


**Supplementary Figure 1. The effect of RhoGEF12 pharmacological inhibition on retromer and SORL1 levels.**

Compared to DMSO control, Y16 results in a small increase in SORL1 in both wildtype mouse cultured neurons (left panel) and in 'wildtype' iNeurons (right panel). (* indicates $p<.05$)

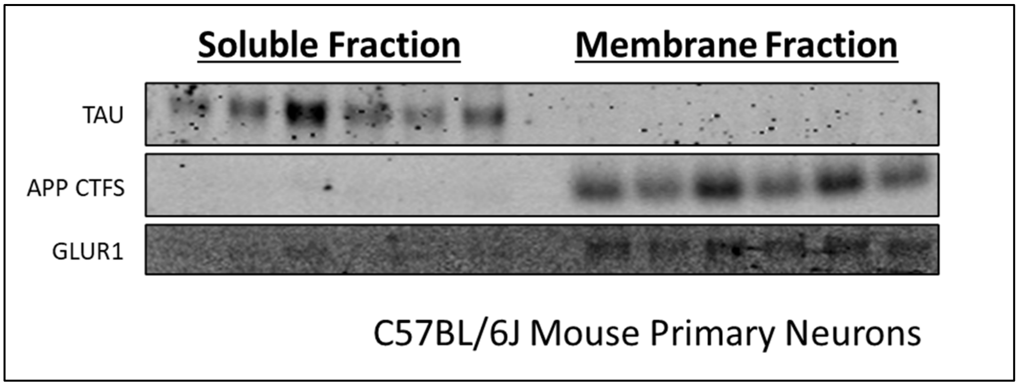


**Supplementary Figure 2. Validating the membrane fractionation protocol**

The soluble fraction is enriched in cytosolic Tau, while the membrane fraction is enriched in endosomal membrane proteins, APP's c-terminal fragment (APP-CTF) and the glutamate receptor GLUR1.
